# Organoid-in-Bead (OrB): vortex-based compartmentalization enables scalable, high-density intestinal organoid culture

**DOI:** 10.64898/2026.06.21.733630

**Authors:** Kazuki Hattori, Hiromi Kirisako, Megumi Matsuo, Sadao Ota

## Abstract

Intestinal organoids are powerful *in vitro* models, but their use in large-scale analyses remains constrained by the low throughput, labor-intensive handling, and high reagent consumption of conventional Matrigel dome culture. Here, we present Organoid-in-Bead (OrB), a vortex-based compartmentalization workflow that partitions organoid fragments into thousands of discrete Matrigel microbeads, enabling scalable, high-density culture from a single batch preparation. OrB maintains dome-comparable organoid growth and epithelial polarity, supports passaging-based culture expansion, yields more than 5,000 organoids in the final 10 cm dish format, and reduces Matrigel and medium consumption by approximately 70% on a per-organoid basis. OrB therefore provides a practical and scalable upstream workflow for generating screening-scale intestinal organoids.

**Highlights:**

- OrB generates Matrigel microcompartments by vortexing without microfluidics
- OrB enables scalable, high-density intestinal organoid culture in one batch
- OrB maintains dome-comparable growth and epithelial polarity and supports passaging
- OrB yields >5,000 organoids per batch with ∼70% less Matrigel/medium per organoid

## Introduction

Large-scale studies using 3D organoid models increasingly depend on the ability to generate sufficient organoid material ahead of downstream assays. Compound screening and genetic perturbation screens in organoids have facilitated unbiased identification of molecular mechanisms that govern tissue function and pathology (Lo et al., 2025; Mead et al., 2022; Ringel et al., 2020; Vlachogiannis et al., 2018). Intestinal organoids are particularly valuable as they recapitulate key features of epithelial barrier function, host-microbe interactions, and therapeutic responses in a physiologically relevant 3D format (Günther et al., 2022).

However, standard Matrigel dome culture remains a practical bottleneck for scaling intestinal organoid experiments (Hofer and Lutolf, 2021). In conventional 30–50 µL Matrigel dome cultures, organoids must be seeded at sufficiently low density to remain spatially separated, preventing aggregation and helping maintain viability (den Daas et al., 2022; Maeda et al., 2023). As a result, output is typically limited to on the order of 100 organoids per dome, and scale-up requires dozens of manually plated domes. This low-density culture format also consumes substantial amounts of Matrigel and medium on a per-organoid basis, significantly increasing reagent burden.

Here, we developed Organoid-in-Bead (OrB), a vortex-based compartmentalization workflow for high-density culture of intestinal organoids (Figure 1). In OrB, brief vortex emulsification partitions organoid fragments into thousands of Matrigel microbeads that serve as microscale culture compartments. This single-batch preparation yields high-density suspension culture of more than 5,000 organoids in the final 10 cm dish, approximately equivalent to the output of 40–50 conventional Matrigel domes, while maintaining physical separation between growing organoids. By converting conventional dome culture into a compartmentalized microscale format, OrB reduces per-organoid Matrigel and culture-medium consumption by approximately 70% and provides an accessible upstream workflow for generating screening-scale intestinal organoid input material.

**Figure 1.**
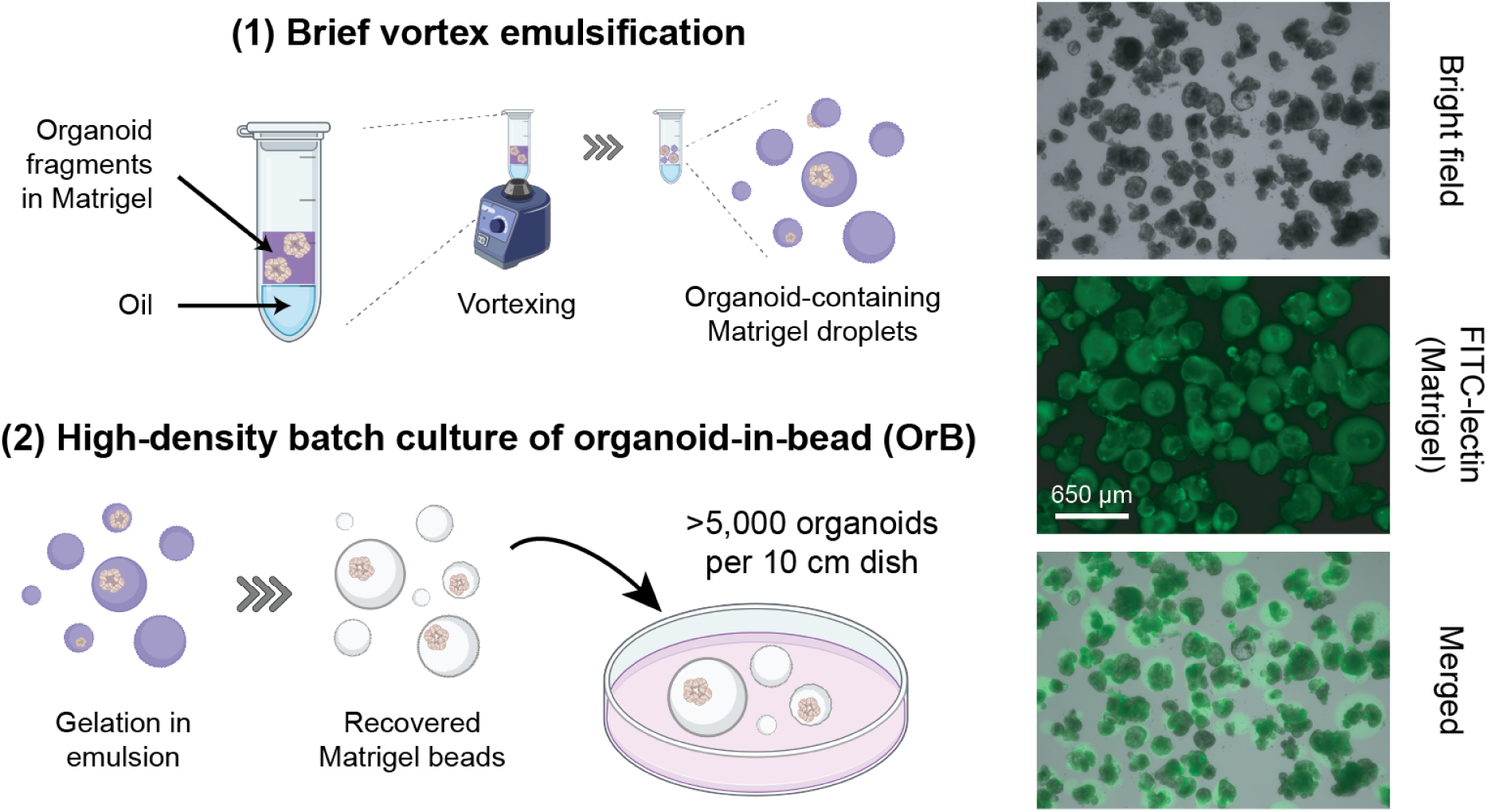
Overview of the Organoid-in-Bead (OrB) workflow. Organoid fragments suspended in 50% Matrigel are layered over an equal volume of surfactant-containing oil and briefly vortexed to generate a water-in-oil emulsion. After gelation of Matrigel at 37°C, the emulsion is broken and Matrigel microbeads are recovered for suspension culture in a dish. This single-batch workflow partitions organoid fragments into thousands of microscale Matrigel compartments and yields more than 5,000 organoids in the final 10 cm dish format by day 5. Representative day 5 images of OrB cultures are shown at right as bright field, FITC-lectin fluorescence, and merged views. Scale bar: 650 µm.

## Results

### Establishment of OrB and benchmarking against conventional dome culture

We first established an OrB workflow to generate Matrigel microbeads that support the growth of mouse intestinal organoids. Using 50% Matrigel layered over 2% Oil for EvaGreen in Noah 7200, a 3-second vortexing step generated water-in-oil droplets that were gelled at 37°C and recovered as Matrigel microbeads. Under this working condition, the recovered microbeads had a median equivalent diameter of approximately 250 µm (Figures 2A and 2B). To define a practical working condition for OrB, we tested a range of oil-phase compositions and vortexing durations and quantified bead sizes from fluorescence images where we stained Matrigel microbeads with a fluorescein isothiocyanate (FITC)-conjugated lectin (Figure 2B). Because our design criterion was to obtain microbeads large enough to allow room for subsequent organoid expansion, we focused on conditions that reproducibly generated microbeads with a median equivalent diameter above 200 µm. Evaluation of multiple conditions showed that bead size decreased with increasing surfactant concentration or longer vortexing duration (Figure 2B). We therefore selected 2% Oil for EvaGreen in Noah 7200 with 3-second vortexing for downstream culture experiments, because this condition reproducibly met the >200 µm size criterion.

**Figure 2.**
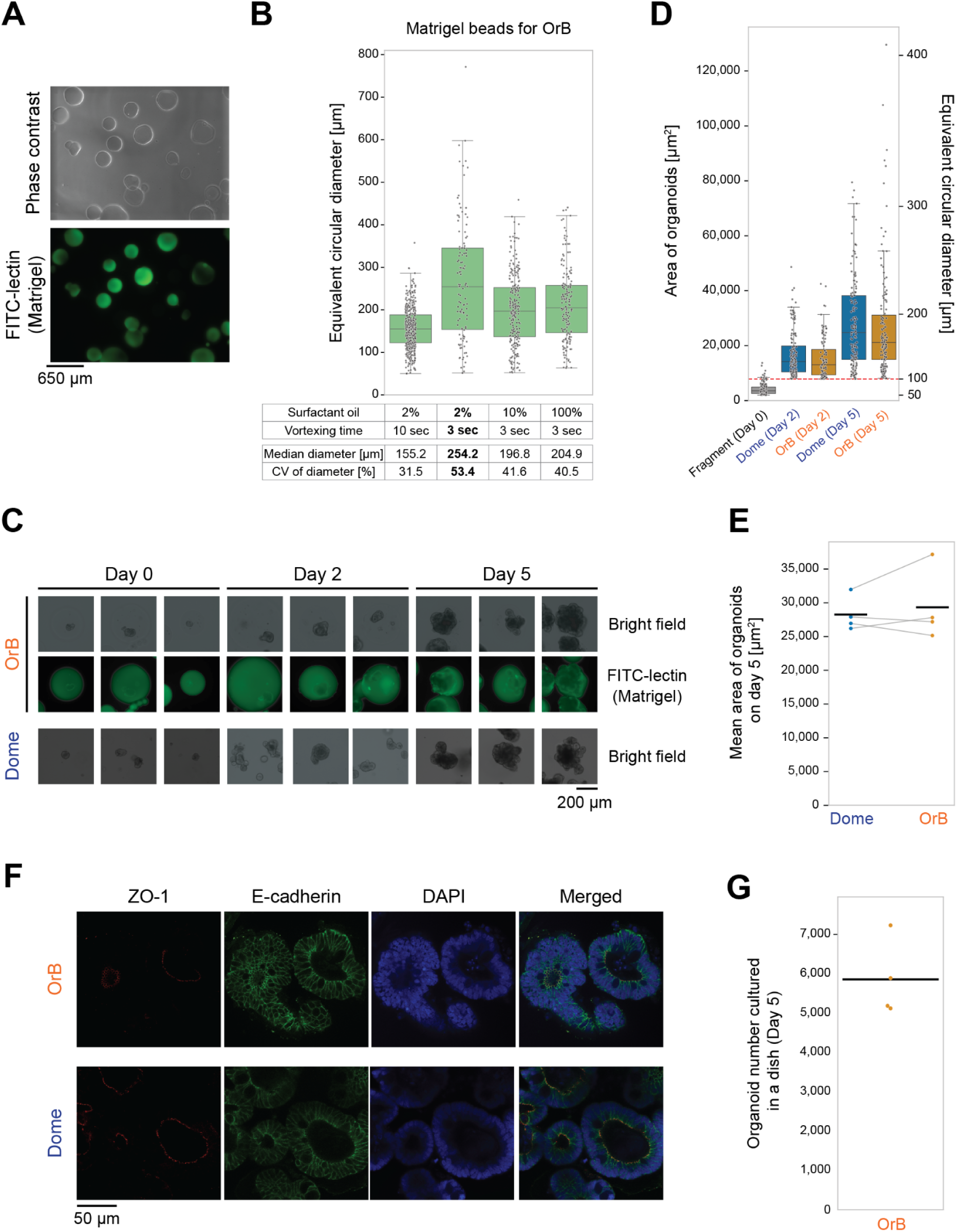
Establishment and benchmarking of OrB for intestinal organoid culture. **(A)** Representative phase contrast and FITC-lectin fluorescence images of Matrigel microbeads generated by the OrB workflow. Scale bar: 650 µm. **(B)** Size distribution of Matrigel microbeads prepared under different surfactant oil compositions and vortexing durations in the OrB workflow. Bead size was quantified from FITC-lectin fluorescence images by image analysis, and expressed as equivalent circular diameter. Box plots with overlaid dots show individual microbeads, and the table summarizes the median diameter and coefficient of variation (CV) for each condition. **(C)** Representative images of parallel OrB and dome cultures generated from the same input organoid fragment preparation and imaged on days 0, 2, and 5. For OrB, bright field images and corresponding FITC-lectin images of Matrigel microbeads are shown; for dome cultures, bright field images are shown. Three representative images are shown for each condition and time point. Scale bar: 200 µm. **(D)** Quantification of organoid growth over time in OrB and dome cultures. Day-0 fragments and organoid on days 2 and 5 were stained with acridine orange (AO), segmented from fluorescence images, and quantified by projected area. Box-and-strip plots show the distribution of individual object areas for a representative experiment; similar trends were reproduced in three additional independent experiments. The red dashed line indicates the organoid size threshold (∼7854 µm^2^, equivalent to a 100-µm-diameter circle) applied to the day 2 and day 5 samples (day 0 fragments, N = 161; dome day 2, N = 158; OrB day 2, N = 104; dome day 5, N = 147; OrB day 5, N = 139). Left y-axis, projected area; right y-axis, equivalent circular diameter. **(E)** Comparison of mean organoid area on day 5. Each dot represents one independent experiment (N = 4), with paired dome and OrB values connected by gray lines. Black horizontal lines indicate the mean. **(F)** Representative confocal immunofluorescence images of day 5 organoids cultured by OrB or by dome culture. Organoids were stained for ZO-1 (tight junction marker), E-cadherin (adherens junction marker), and DAPI nuclear counterstaining. Scale bar: 50 µm. **(G)** Estimated total organoid yield of OrB cultures on day 5. Organoid concentration was determined from AO-stained aliquots imaged in a defined volume and multiplied by the total suspension volume. Each dot represents one independent experiment (N = 4), and the black line indicates the mean. The measured yield corresponded to more than 5,000 organoids per batch in the final 10 cm dish format.

To benchmark OrB against conventional dome culture for culturing mouse small intestinal organoids, we performed a side-by-side comparison using the same input organoid fragments. We first mechanically fragmented dome-grown organoids through vigorous pipetting, and split these fragments in parallel into OrB microbeads and new domes. During culture, we exchanged culture medium on days 2 and 4 in both workflows. Representative bright field and FITC-lectin images showed similar expansion of organoid structures in OrB and dome cultures on days 2 and 5 (Figure 2C).

To quantify organoid growth, we stained organoids with AO to label the entire structure, acquired 2D fluorescence images, and measured projected area by image segmentation. Before imaging, organoids were recovered from domes by gentle mechanical disruption and from OrB cultures using a Matrigel dissociation reagent. We defined organoids as AO-positive objects with an area at least equivalent to a 100-µm-diameter circle (∼7854 µm^2^), in line with size criteria used in prior organoid studies (Lee et al., 2021; Yang et al., 2021); across four independent experiments, this threshold corresponded to the 88.14 ± 4.60 percentile of the day 0 fragment size distribution. OrB and dome cultures showed similar growth trajectories over 5 days (Figure 2D). Across four independent experiments, mean organoid area on day 5 was comparable between conditions (Figure 2E), indicating that OrB supports organoid growth comparable to conventional dome culture under the tested condition.

We further characterized epithelial polarity in day 5 OrB-derived organoids by immunofluorescence staining and confocal microscopy. OrB-derived organoids exhibited membrane-localized E-cadherin and apically localized ZO-1, indicating the formation of polarized epithelial structures with an apical-in organization. Similar staining patterns were observed in dome-derived organoids (Figure 2F).

Next, we quantified the culture scale achievable with OrB under the tested batch workflow by measuring organoid yield on day 5. Using the organoid definition described above, we imaged all organoids within a defined volume of the OrB suspension. We then calculated organoid concentration and estimated total yield by multiplying by total suspension volume. Across four independent experiments, the tested OrB workflow yielded 5,760 ± 905 organoids per batch by day 5 (mean ± SD). Thus, OrB supports a high-density culture of more than 5,000 mouse intestinal organoids in the final 10 cm dish (Figure 2G).

To assess the reagent efficiency, we compared per-organoid Matrigel and medium consumption between dome culture in a 24-well plate and OrB culture in the tested batch workflow (Table 1). In the experiment, we exchanged medium on days 2 and 4 and counted organoids on day 5 by AO staining and image analysis. On a per-organoid basis, OrB reduced Matrigel consumption from 0.1521 to 0.0427 µL (71.9% reduction) and reduced medium consumption from 11.41 to 3.42 µL (70.0% reduction). Together, these results indicate that OrB reduces per-organoid Matrigel and medium consumption by about 70%.

**Table 1.**
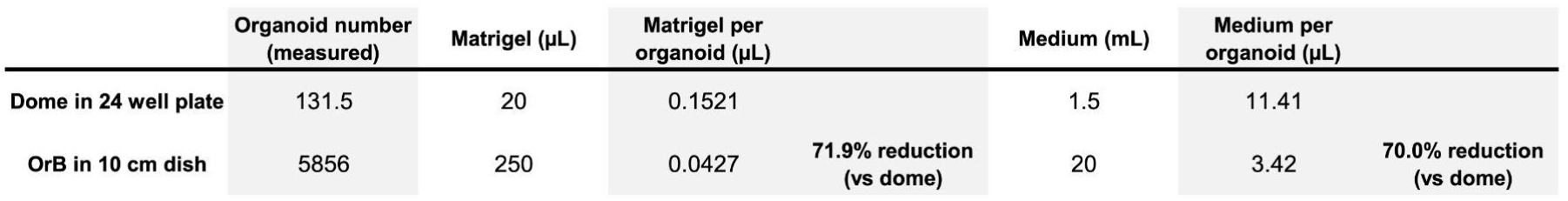
Per-organoid Matrigel and medium consumption in OrB and conventional dome culture.

To assess reagent efficiency, we compared per-organoid consumption of Matrigel and culture medium between conventional dome culture in a 24-well plate and OrB culture, which used a 60 mm dish from days 0 to 2 and a 10 cm dish thereafter. Organoids were cultured with medium changes on days 2 and 4 and counted on day 5 by AO staining and image analysis.

For yield calculation, only AO-positive objects with a projected area at least equivalent to a 100-µm-diameter circle (∼7854 µm^2^) were counted as organoids. The table reports total undiluted Matrigel and culture medium volumes used, per-organoid consumption, and the percent reduction in OrB relative to dome culture.

### OrB supports organoid expansion by single-batch passaging

To enable expansion of OrB cultures, we established a batch-format passaging workflow for organoids grown in OrB (Figure 3A). We recovered mature organoids (days 4-6) by Matrigel dissolution, fragmented them by passing through a 70-µm mesh, and quantified them by AO-based image analysis to normalize seeding density (10,000 fragments per 500 µL of 50% Matrigel). The resulting fragments were then re-encapsulated into OrB, cultured in a 60 mm dish, and transferred to a 10 cm dish on day 2.

**Figure 3.**
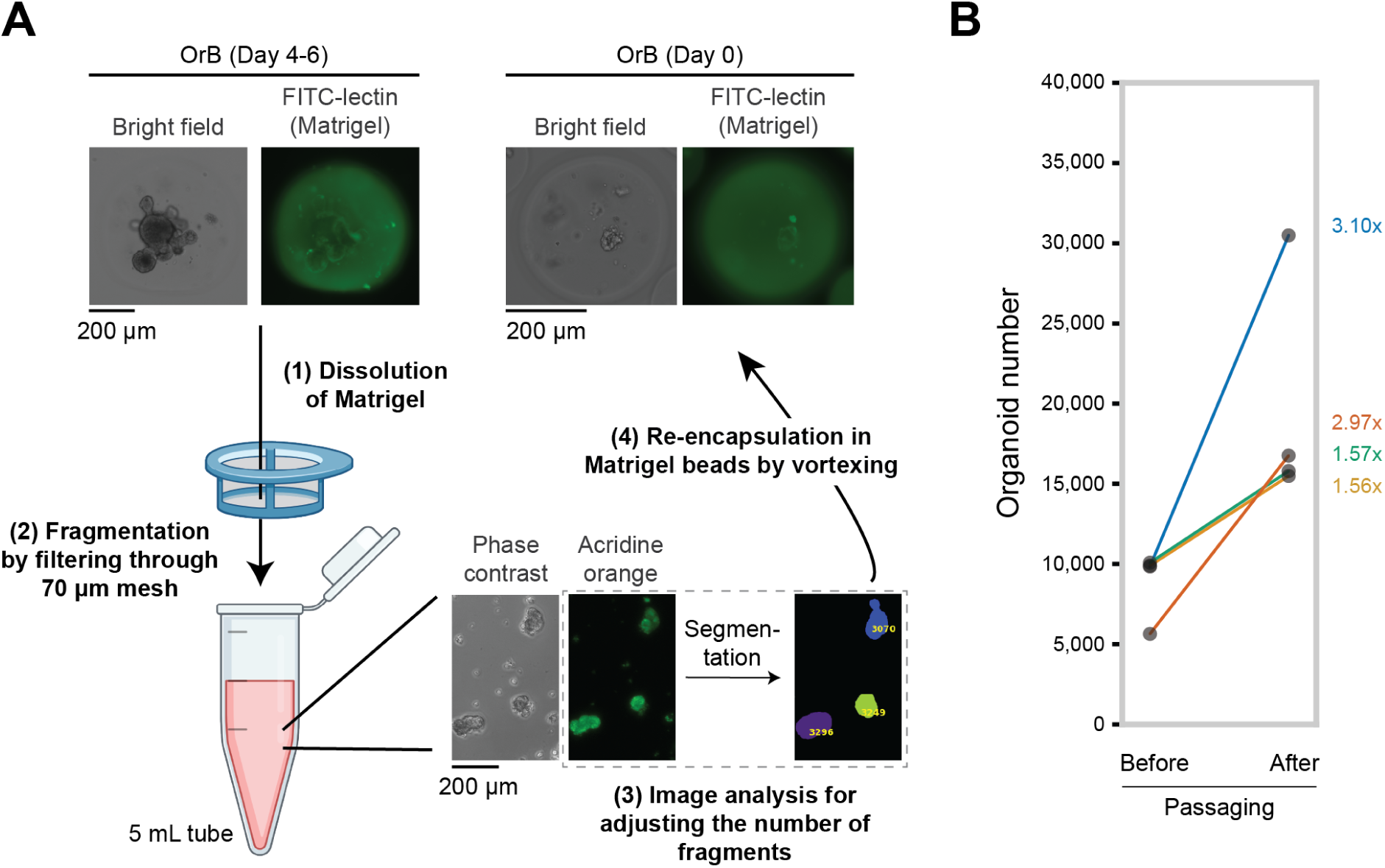
Batch passaging of OrB cultures supports organoid expansion. **(A)** Workflow for OrB-based organoid passaging and expansion. Mature organoids cultured by OrB were recovered by Matrigel dissolution, fragmented through a 70 µm mesh, quantified by AO-based image analysis, and re-encapsulated into OrB at a defined input density. A subset of 10,000 fragments was used to produce new OrB, cultured in a 60 mm dish for 2 days, and subsequently scaled up to a 10 cm dish for an additional 2 to 4 days to regenerate organoids. **(B)** Paired comparison of organoid numbers before passaging and estimated total organoid yield after OrB passaging. Each line represents an independent experiment (N = 4). Because only a subset of recovered fragments was reseeded, after-passaging values were estimated by scaling the measured organoid number in the cultured subset by the ratio of total fragments collected to fragments seeded. Fold changes are shown to the right of the graph, with text colors matching the corresponding line colors.

To assess whether this passaging workflow expands organoid cultures, we compared organoid numbers by image analysis immediately before passaging and with the estimated total organoid output 4–6 days after re-encapsulation, using the same workflow described in Figure 2G. After passaging, we seeded only a subset of the collected fragments into the next OrB culture, and estimated the total post-passaging yield by multiplying the organoid count measured in the cultured subset by the ratio of total fragments collected to fragments seeded (total yield = total organoids measured × total fragments collected / fragments seeded).

Across four independent experiments, OrB passaging produced an estimated 1.56 to 3.10-fold increase in organoid numbers (Figure 3B). These results indicate that OrB culture can be passaged in batch format and can regenerate organoid populations on the order of ten thousand organoids per passage.

### OrB supports organoid formation directly from primary crypts

To test whether OrB can generate organoids directly from primary tissue, we applied the OrB workflow to freshly isolated mouse small intestinal crypts (Figure 4A). The encapsulated crypts formed organoids over culture, as confirmed by bright field microscopy. To normalize organoid output to crypt input, we quantified crypt number before encapsulation by staining with a phycoerythrin (PE)-conjugated anti-CD44 antibody and image analysis, taking advantage of preferential CD44 labeling of crypt epithelial regions relative to villus epithelium (Wang et al., 2013).

**Figure 4.**
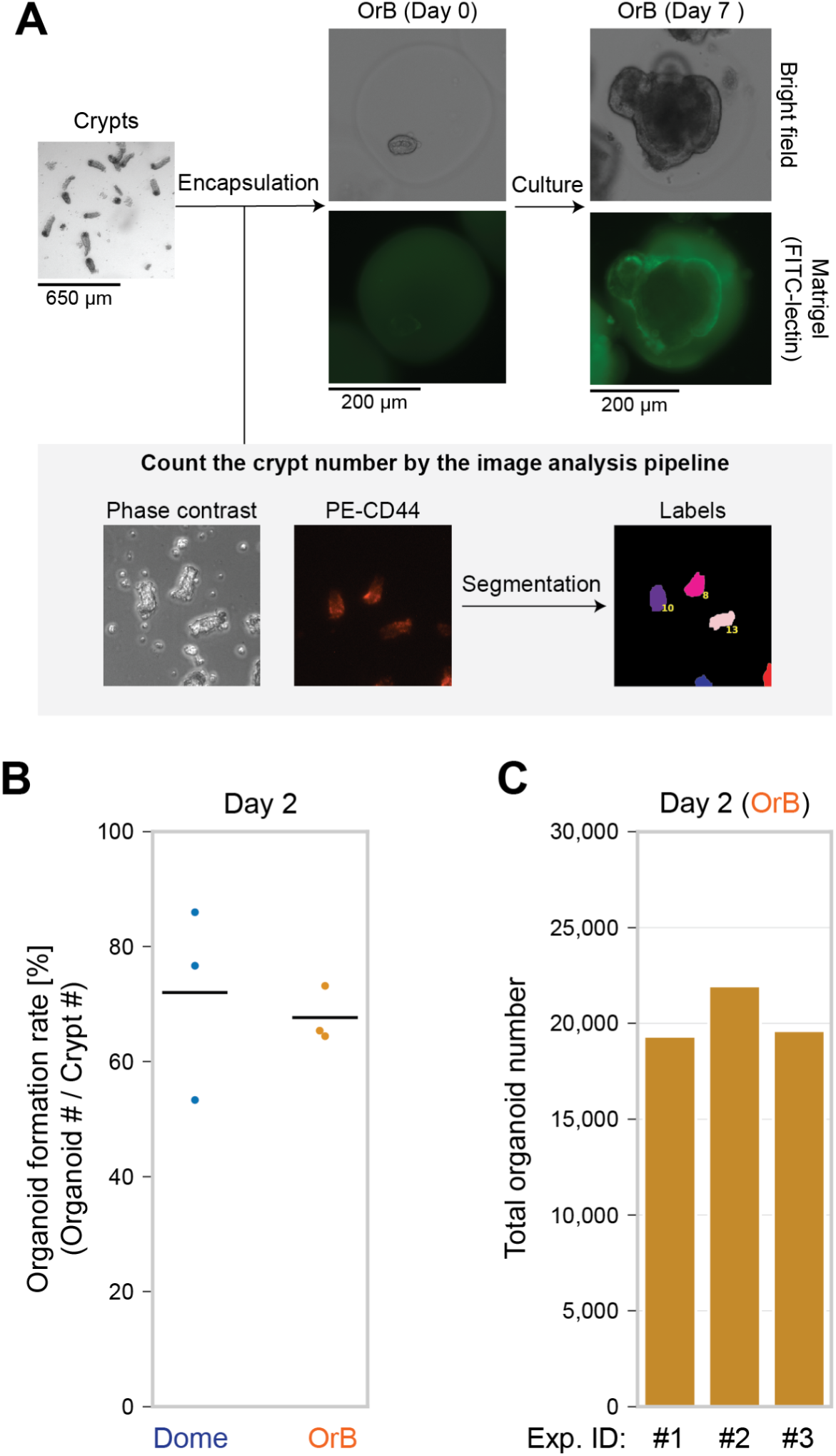
Organoid formation from crypts by OrB. **(A)** Workflow schematic for OrB-based organoid generation and crypt counting. Isolated mouse small intestinal crypts were stained with PE-conjugated anti-CD44 antibody, which preferentially labels crypt regions but not villi, and input crypt number was quantified by automated image analysis. Crypts were then encapsulated in Matrigel microbeads (OrB) and cultured to form organoids. Representative images of OrB cultures are shown on days 0 and 7. Matrigel was visualized by FITC-lectin staining. Scale bars: 650 µm for low-magnification images and 200 µm for enlarged images. **(B)** Organoid formation rate in dome and OrB cultures on day 2, calculated as the number of day-2 organoids divided by input crypt number. Both input crypt numbers and day-2 organoid numbers were quantified by image analysis. Each dot represents one independent experiment (N = 3), and black lines indicate the mean. **(C)** Total organoid yield in OrB cultures initiated with 30,000 crypts. Bars represent three independent experiments using crypts isolated from three different mice. Organoids were quantified on day 2 by AO staining and image analysis.

We next compared the efficiency of organoid generation from primary crypts between OrB and conventional dome culture. Because crypt-derived organoids are typically small on day 2, we defined day-2 organoids as AO-positive objects with a projected area at least equivalent to a 50-µm-diameter circle (∼1,963.5 µm^2^) and calculated the formation rate by dividing the day-2 organoid count by the CD44-based input crypt count before encapsulation. Under the tested conditions, OrB achieved a comparable formation rate to dome culture on day 2 (dome, 72.00 ± 16.83%; OrB, 67.66 ± 4.80%; mean ± SD, N = 3; Figure 4B). OrB culture initiated with 30,000 crypts yielded approximately 20,000 organoids per batch by day 2 across three independent experiments from three different mice (Figure 4C). Together, the results show that OrB can generate screening-scale intestinal organoid material directly from primary crypts while maintaining formation rates similar in magnitude to those of dome culture.

## Discussion

We developed OrB, a vortex-based Matrigel microbead culture that enables screening-scale production of intestinal organoids while reducing per-organoid Matrigel and medium consumption by more than 70%. OrB supported dome-comparable organoid growth and polarized epithelial architecture, yielded approximately 5,000 organoids in the final 10 cm dish, and enabled batch passaging with 1.56 to 3.10-fold expansion. OrB also generated organoids directly from primary crypts, producing approximately 20,000 organoids per batch by day 2 while maintaining formation rates comparable to those observed in dome culture.

Together, these results establish OrB as a practical upstream method for preparing screening-scale intestinal organoid input material from two intestinal starting materials, organoid fragments and primary crypts.

The design principle of OrB is microscale compartmentalization with standard laboratory instrumentation. Suspension culture at low Matrigel concentration avoids solid-gel embedding and simplifies handling and automation; however, bulk suspension expansion can alter epithelial polarity in organoids (Hirokawa et al., 2021). Suspended microliter-scale hydrogels improve scalability by moving larger matrix units into suspension, but do not increase organoid density within each matrix unit (Co et al., 2023). Microfluidic encapsulation can generate highly uniform hydrogel microbeads containing organoids, but it requires specialized devices and flow control (Lavickova et al., 2026; Maekawa et al., 2026). In contrast, OrB generates thousands of discrete Matrigel compartments using only a vortex mixer, thereby converting conventional dome culture into a dense batch format that maintains physical separation between growing organoids while preserving dome-comparable growth and polarized epithelial architecture.

OrB also has important limitations. Because vortex emulsification is stochastic, neither bead size nor the number of organoid fragments per bead can be precisely specified. To prevent within-bead merging, organoids must be seeded at low occupancy, which produces many empty microbeads and reduces reagent efficiency relative to an ideal one-organoid-per-bead system. In addition, the scale demonstrated here, 5,000 organoids per batch, reflects batch production from a single 2 mL emulsification tube, and further scale-up could be pursued by multiplying tubes in parallel and/or adapting the workflow to larger tube formats such as 50 mL tubes.

Finally, we validated OrB only in mouse small-intestinal organoids, using organoid fragments and primary crypts as starting materials. The extent to which this workflow can be adapted to other Matrigel-dependent organoid systems remains to be determined. Even with these constraints, OrB addresses a practical bottleneck in organoid workflows: the upstream generation of large numbers of physically separated organoid units. We view OrB not as a universal replacement for dome culture, but as an accessible manufacturing format for generating dome-like intestinal organoid input material at high density and in batch format.

Beyond this upstream role, OrB may also interface naturally with downstream high-throughput microcompartment-based phenotypic assays (Hattori, 2026), such as 3D imaging flow cytometry (Yamashita et al., 2024) and multimodal analysis (Kawasaki et al., 2024).

## Methods

### Preparation of imaging chambers for organoids and crypts

We assembled 0.43 mm-height imaging chambers using 0.43 mm-thick double-sided tape (NICHIBAN, NW-N30). We first laminated the tape onto a release paper sheet (150 mm × 105 mm; Kyowa Shiko) and cut it into four strips (approximately 30 mm × 3 mm). We then affixed the four strips to the center of a 24 × 60 mm coverslip (Matsunami Glass, C024601) to form a rectangular frame, placing the two long strips along the long axis of the coverslip and closing the rectangle by attaching the short strips tightly to the ends of the long strips. Excess tape extending beyond the coverslip edges was trimmed with scissors. For easier handling and imaging, we affixed the tape-framed coverslip to a microscope slide glass (Matsunami Glass, S2112) using masking tape. Immediately before sample loading, we removed the top release liner from the tape frame, dispensed the sample into the central cavity while avoiding air bubbles, and sealed the chamber by gently placing a second coverslip (24 × 40 mm; Matsunami Glass, C024401) on top to ensure full contact with the tape surface. Because organoids readily adhere to the bottom coverslip, we promptly placed the top coverslip to distribute the organoids evenly across the chamber before they adhered.

### Isolation of mouse intestinal crypts

The isolation procedure was based on a previously published protocol, with minor modifications (Imada et al., 2024). We collected the distal small intestine (approximately the distal half) from 8-week-old male C57BL/6JmsSlc mice (Japan SLC, Inc.) immediately after euthanasia and transferred it to ice-cold D-PBS(-) (045-29795, Fujifilm Wako). We removed mesentery and visible fat, opened the intestine longitudinally, and gently rubbed the luminal surface between fingers to remove mucus.

We washed the opened intestine in a 10 cm petri dish or 6-well plate containing ice-cold D-PBS(-), cut it into 4 to 5 pieces, and transferred the pieces to a 50 mL tube containing 30 mL of ice-cold D-PBS(-) supplemented with 10 mM EDTA (0.5 M, pH 8.0; Thermo Fisher, AM9260G). We rocked the tube for 40 min on a rotating shaker (NISSIN, NRC-30D) at 4°C to dislodge epithelial crypts.

After EDTA treatment, we gently shook the tissue by hand 50 times. We moved the intestinal fragments to a well of a 6-well plate containing 2 mL of ice-cold D-PBS(-) and washed them by gentle agitation. We removed excess liquid with a paper towel and transferred the tissue pieces to a new 50 mL tube. We then added 20 mL of ice-cold D-PBS(-) and shook the tube vigorously 70 times to release crypts.

We poured the supernatant containing released crypts through a 70 µm nylon mesh folded into four layers (AsOne, 62-0866-44) shaped into a funnel, collecting the flow-through as the first fraction. We repeated this shaking and filtration step 1 to 2 more times to obtain the second and third fractions, which we collected in separate 50 mL tubes.

We transferred 200 µL of each fraction to wells of a 48-well plate and acquired bright field images on an EVOS M7000 imaging system (Thermo Fisher) using an automated acquisition protocol to confirm villus contamination. Based on these images, we selected fractions enriched for crypts for further processing.

We pooled crypt-enriched fractions as needed and centrifuged them at 300 × g for 5 min at 4°C. When crypts failed to form a visible pellet, we repeated the centrifugation step. We discarded the supernatant, resuspended the pellet in 10 mL of ice-cold intestinal stem cell basal medium (ISC basal medium; Advanced DMEM/F-12 (Thermo Fisher Scientific, 12634010) supplemented with 1x L-alanyl-L-glutamine (100x; Fujifilm Wako, 016-21841) and 1x HEPES (100x; Fujifilm Wako, 345-06681)), and transferred the suspension to a 15 mL tube. We centrifuged the suspension at 200 × g for 3 min at 4°C, removed the supernatant, and resuspended the pellet in 10 mL of ice-cold ISC basal medium. To remove single cells, we centrifuged the suspension at 100 × g for 1 min at 4°C. We discarded the supernatant and resuspended the crypt pellet in approximately 4 mL of ice-cold ISC basal medium, adjusting the volume to match the pellet size.

The Animal Care and Use Committee of the Research Center for Advanced Science and Technology (RCAST), The University of Tokyo, approved all animal experiments.

### Crypt counting through CD44 staining and imaging

To count crypts in a semi-automated and reproducible manner, we stained crypts with an anti-CD44 antibody, which preferentially stains crypt regions rather than villi. For the staining, we transferred 100 µL of the crypt suspension to a 1.5 mL ProteoSave low-binding tube (Sbio, MS-4215M) using a 200 µL wide-bore pipette tip. We marked the 100 µL liquid level on the tube wall to standardize supernatant removal in later steps. We added 1 µL of PE-conjugated anti-mouse CD44 antibodies (BioLegend, 103023), mixed gently, and incubated the tube at 37°C in a 5% CO2 incubator for 10 min.

After incubation, we added 1 mL of D-PBS(-) containing 0.1% Poloxamer 188 (Thermo Fisher, 24040-032) and mixed the suspension by pipetting with a standard 1,000 µL tip to disperse the crypts. We briefly centrifuged the tube in a mini centrifuge for about 10 s and removed the supernatant down to the 100 µL mark.

We resuspended crypts in 500 µL of D-PBS(-) containing 0.1% Poloxamer 188. We then loaded 100 µL of the suspension into the custom-made imaging chamber using a 200 µL-wide-bore tip. We acquired tiled phase contrast and fluorescence images of PE-CD44 (EVOS Light Cube, RFP 2.0 (AMEP4952)) covering the entire chamber area on an EVOS M7000 imaging system using an automated acquisition protocol and saved all images as raw TIFF files.

We analyzed the images with a custom Python pipeline to count crypts in 100 µL of suspension. The script segmented crypt-like structures and quantified objects that met predefined size criteria to estimate crypt counts and crypt concentration, as described in the image analysis section.

### Routine maintenance culture of mouse small intestinal organoids

All organoids used in this study were derived from mouse small intestinal crypts. Except for experiments performed in OrB, we maintained organoids in conventional Matrigel (Corning, 356231) domes. We seeded seven domes per well in a 6-well plate, using 40 µL of 50% Matrigel per dome, and cultured the organoids in 2 mL of ENR medium (see details below) per well. For routine maintenance, the medium was changed every 2 days. In experiments directly comparing OrB and dome cultures, the medium was changed on days 2 and 4.

To prepare ENR medium, we first prepared ISC crypt medium, which consisted of ISC basal medium supplemented with 1x B-27 Supplement (50x; Thermo Fisher, 17504044), 1x N2 Supplement with Transferrin (Holo) (100x; Fujifilm Wako, 141-08941), 1 mM N-acetyl-L-cysteine (Fujifilm Wako, 017-05131), and 1% Antibiotic-Antimycotic (Thermo Fisher, 15240096). We then supplemented the ISC crypt medium with 50 ng/mL mouse EGF (Miltenyi Biotec, 130-094-037), 100 ng/mL mouse Noggin (Miltenyi Biotec, 130-103-459), and 20% (v/v) conditioned medium from Cultrex HA-R-Spondin1-Fc 293T cells (R&D Systems, 3710-001-01) as the source of R-spondin 1.

### Dome culture conditions for benchmarking against OrB

For side-by-side benchmarking against OrB, conventional dome cultures were prepared in 24-well plates. Each well contained one Matrigel dome, prepared using 40 µL of 50% Matrigel containing organoid fragments, and was cultured in 500 µL of ENR medium.

Medium was replaced with 500 µL of fresh ENR medium on days 2 and 4. Organoids were counted on day 5 by AO staining and image analysis as described below.

### Generation and culture of OrB

#### Preparation of organoid fragments for OrB encapsulation

We collected organoids grown in Matrigel domes from a 6-well plate into a 5 mL microcentrifuge tube (Eppendorf, 0030119401). To minimize organoid adhesion, we pretreated tubes and pipette tips with CELLOTION (Zenogen, 11918) before use. We disrupted Matrigel by vigorously pipetting 30 times with a P1000 pipette set to 1,000 µL, then pelleted the organoids at 200 × g for 3 min. When we visually confirmed the presence of a residual gel layer, we resuspended the pellet in 1 to 5 mL of ice-cold ISC basal medium and then repeated centrifugation at 200 × g for 3 min. We discarded the supernatant and resuspended the pellet in 200 µL of ISC basal medium.

We mechanically fragmented organoids by vigorous pipetting 200 times with a P200 pipette set to 200 µL. We then added ISC basal medium to a final volume of 4 mL without mixing, then pelleted the fragments at 200 × g for 3 min. We discarded the supernatant, resuspended the pellet in 4 mL of ISC basal medium, and dispersed fragments by pipetting approximately 10 times with a P1000 pipette. To remove single cells, we centrifuged the suspension at 100 × g for 1 min, discarded the supernatant, and resuspended the fragment pellet in 1 mL of ISC basal medium.

### Imaging-based quality control of organoid fragment concentration

To measure fragment size distribution and concentration, we mixed 10 µL of the fragment suspension with 15 µL of ISC basal medium and 25 µL of 0.001% AO (Fujifilm Wako, 014-08941) in D-PBS(-). We loaded the entire mixture into the custom-made imaging chamber. We acquired tiled images using an EVOS M7000 imaging system with a 4x objective in phase contrast and GFP channels (EVOS Light Cube GFP 2.0, Thermo Fisher, AMEP4951). To maintain consistent acquisition parameters across experiments, we loaded the imaging settings file from the prior experiment and used the same acquisition template.

We saved tiled images as raw TIFF files for both channels and used a stitch setting. When fragments were too dense for reliable segmentation, we diluted the suspension by increasing the volume of the ISC basal medium and repeating imaging.

### Preparation of organoid fragment suspension for OrB generation

We adjusted the fragment suspension to 10,000 fragments per 250 µL, aliquoted 250 µL into a 1.5 mL tube, and chilled it on ice for 5 min. We maintained this fragment concentration because fewer fragments can reduce cell density and impair regrowth, whereas more fragments increase the fraction of microbeads containing multiple fragments and can promote aggregation-induced cell death. To prevent gelation of Matrigel, we kept the samples at 4°C until vortexing.

We added 250 µL of chilled Matrigel to the 250 µL of fragment suspension and mixed by pipetting 30 times with a P200 pipette to minimize fragment aggregation. We layered the resulting 500 µL suspension onto 500 µL of prechilled oil phase (2% (v/v) Oil for EvaGreen (Bio-Rad, 1864112) in Noah 7200 (Zhejiang NOAH Fluorochemical)) in a 2 mL tube.

### Emulsification, gelation, emulsion breaking, and culture

To emulsify, we vortexed the tube for 3 seconds using a vortex mixer (Scientific Industries, Digital VORTEX-GENIE 2) at maximum speed (3,200 rpm). We visually confirmed the absence of residual bulk aqueous phase after mixing. We gelled Matrigel by incubating the tubes on a Thermoblock Rotator (NISSIN, SN-48BN) at 37°C (speed setting "2") for 20 min. After gelation, we removed the bottom oil phase beneath the droplet layer using a syringe with a needle. We then overlaid 500 µL of ENR medium and added 500 µL of Noah 7200 containing 20% (v/v) 1H,1H,2H,2H-perfluoro-1-octanol (PFO; Fujifilm Wako, 324-90642) to the bottom of the tube to initiate demulsification. We inverted the tube while applying a plasma ball (Y-Depart Center 55) to induce electrocoalescence and demulsification.

Because emulsion breaking was often incomplete in a single step, we recovered the aqueous ENR phase in multiple rounds. Using a CELLOTION-pretreated 1,000 µL wide-bore tip, we transferred the recovered ENR phase to a 60 mm cell culture dish, added a fresh 500 µL of ENR medium to the tube, and repeated plasma ball-assisted breaking. We repeated this cycle four times to recover a total of 2 mL of ENR medium in four 500 µL fractions. When droplets were difficult to break, we replaced the 20% (v/v) PFO in Noah 7200 with a freshly prepared solution.

To recover residual Matrigel microbeads, we transferred the remaining aqueous phase, containing a small amount of oil, into a new 1.5 mL tube, allowed the phases to separate for several seconds, and collected the ENR layer into the same 60 mm cell culture dish. Finally, we adjusted the culture volume to 4 mL with ENR medium, mixed gently, and initiated culture in a 5% CO_2_ incubator.

On day 2, we refreshed the culture medium and transferred OrB to a 10 cm cell culture dish. We collected the OrB suspension from the 60 mm dish using a cell lifter, transferred it to a 15 mL tube, centrifuged at 100 × g for 3 min, and removed the supernatant. We resuspended the OrB pellet in 8 mL of fresh ENR medium and transferred it to a 10 cm dish. On day 4, we collected the OrB suspension, centrifuged it at 100 × g for 3 min, replaced the supernatant with 8 mL of fresh ENR medium, and returned the culture to the original 10 cm dish.

### Passaging of OrB

#### Collection of OrB and Matrigel dissolution

We collected OrB cultures from 10 cm dishes by first dislodging microbeads attached to the dish bottom with a cell lifter. We transferred the bulk OrB suspension into a 15 mL STEMFULL tube (Sbio, MS-90150) using a 5 mL disposable pipette, then tilted the dish and used the cell lifter again to recover remaining OrBs. We rinsed the dish with 2 to 4 mL of ISC basal medium and then transferred the rinse to the same 15 mL tube using a 5 mL disposable pipette. We did not rewash the dish with the bead suspension because this step caused the microbeads to reattach to the dish, thereby reducing recovery. We pelleted microbeads at 200 × g for 3 min and discarded the supernatant.

To dissolve Matrigel, we added 4 mL of ice-cold Cell Recovery Solution (Corning, 354253) to the pellet using a 5 mL disposable pipette, without mixing. As a guideline, we used 2 mL of Cell Recovery Solution per approximately 500 µL of gel pellet, increasing the volume as the gel pellet volume increased. We secured the tube on a rotary mixer (NISSIN, NRC-30D) placed in a 4°C refrigerator and rotated at the maximum speed (15 rpm) for 20 min. To fully disperse the pellet, we slowly pipetted the suspension ∼30 times using an ice-cold CELLOTION-pretreated P1000 tip before rotation.

After the incubation, we added 10 mL of ISC basal medium and centrifuged at 200 × g for 3 min. Because gel remnants often floated at the liquid surface, we first aspirated and retained ∼2 mL of the upper fraction using a pipette. We then removed the remaining supernatant, combined the upper fraction with the retained fraction, and gently pipetted ∼30 times with an ice-cold CELLOTION-pretreated P1000 tip to disperse residual gel. We added 10 mL of ISC basal medium and centrifuged again at 200 × g for 3 min. We discarded the supernatant and resuspended the pellet in 3 mL of ISC basal medium using a 5 mL disposable pipette, then gently dispersed the pellet with an ice-cold CELLOTION-pretreated P1000 tip until homogeneous.

### Preparation of fragments for passaging

For passaging, we first dispersed organoid aggregates in the 15 mL tube by pipetting 40 times using a CELLOTION-pretreated P1000 tip. We then generated fragments by forcing the suspension through a 70 µm mesh into a 5 mL tube while keeping the pipette tip pressed tightly against the mesh. We rinsed the 15 mL tube with 1 mL of ISC basal medium, passed the rinse through the same mesh to recover any remaining organoids, and repeated this step once more.

We pelleted fragments at 200 × g for 3 min, discarded the supernatant, and resuspended the pellet in 5 mL of ISC basal medium. We pipetted 30 times using a CELLOTION-pretreated P1000 tip. We centrifuged again at 200 × g for 3 min, discarded the supernatant, resuspended the pellet in 2 mL of ISC basal medium, and stored the suspension at 4°C until encapsulation.

We adjusted fragment concentration by staining fragments with AO as described in the “Imaging-based quality control of organoid fragment concentration” section and quantifying them by image analysis as described in the “Image analysis” section. We then generated and cultured OrB as described in the “Preparation of organoid fragment suspension for OrB generation” and “Emulsification, gelation, emulsion breaking, and culture” sections.

### FITC-lectin staining and imaging of Matrigel microbeads

We aliquoted 180 µL of the OrB into a 1.5 mL ProteoSave low-binding tube using a 200 µL wide-bore pipette tip. We added 20 µL of 1 mg/mL FITC-conjugated lectin (from Triticum vulgaris; Sigma, L4895-2MG) in D-PBS(-) to a final concentration of 50 µg/mL, and mixed gently while minimizing contact with the tube wall to reduce bead loss due to adhesion. We incubated the tube at 37°C for 30 min without agitation.

After staining, we added 1 mL of FluoroBrite DMEM (Thermo Fisher, A1896701) and centrifuged the suspension at 200 × g for 3 min. We discarded the supernatant, resuspended the pellet in 1 mL of FluoroBrite DMEM, and centrifuged again at 200 × g for 3 min. We removed the supernatant, leaving approximately 200 µL to avoid disturbing the pellet and maintain bead recovery.

We loaded 200 µL of the stained bead suspension into the custom-made imaging chamber and acquired images on an EVOS M7000 imaging system using a 4x objective in phase contrast and the FITC fluorescence channel (EVOS Light Cube GFP 2.0).

### Epifluorescence imaging for the organoid area and number

We pre-aliquoted 1 mL of D-PBS(-) containing 0.1% (w/v) Poloxamer 188 into a 1.5 mL tube, added 100 µL of the organoid suspension using a CELLOTION-pretreated wide-bore tip, and centrifuged at 200 × g for 3 min. We discarded the supernatant and resuspended the pellet in 200 µL of D-PBS(-) containing 0.1% (w/v) Poloxamer 188. We then added 200 µL of 0.001% (w/v) AO in D-PBS(-), mixed gently, and loaded 200 µL into the custom-made imaging chamber.

For organoid counting, we acquired tiled images on an EVOS M7000 microscope using a 4x objective in phase contrast and the AO fluorescence channel (EVOS Light Cube GFP 2.0) and saved stitched tiled raw images as TIFF files. When organoid density was too high for reliable segmentation, we diluted the sample with 0.0005% (w/v) AO and reimaged it. For organoid area measurements, we acquired non-tiled field images under the same staining and imaging conditions.

### Immunocytochemistry using 3D confocal microscopy

The procedure was based on a previously published protocol, with minor modifications (Martinez-Ordoñez et al., 2023). We prepared Organoid Wash Buffer (OWB) to a final volume of 250 mL by mixing 25 mL of 10x D-PBS(-) (final 1x, Fujifilm Wako, 163-25265), 0.5 g bovine serum albumin (BSA; final 0.2% w/v, Fujifilm Wako, 015-27053), 0.125 mL of 20% SDS (final 0.01% (w/v), Nippon Gene, 311-90271), and 2.5 mL of 10% Triton X-100 (final 0.1% (v/v), Sigma, QJ-T8787-100ML), then bringing the volume to 250 mL with MilliQ water (222.3 mL). We stored OWB at 4°C for up to 1 month. We prepared Phosphate Buffered Saline with Tween 20 (PBST) to a final volume of 1 L by mixing 100 mL of 10x

D-PBS(-) and 1 mL Tween-20 (final 0.1% (v/v), Sigma, MSAP1379-100ML), then bringing the volume to 1 L with MilliQ (899 mL). We stored PBST at room temperature (approximately 22°C to 24°C) for up to several months. We used wide-bore tips for all pipetting steps involving organoids, and we coated all materials that contacted organoids with CELLOTION before use.

Day 0: We collected organoids in culture medium by mechanically disrupting Matrigel domes or microbeads through pipetting, then centrifuged the suspension at 200 × g for 3 min at 4°C. We removed the supernatant, washed the organoids with 1 mL of D-PBS(-), and centrifuged again at 200 × g for 3 min at 4°C. After removing D-PBS(-), we fixed organoids by immersing them in 1 mL of ice-cold 4% paraformaldehyde in D-PBS(-) (Fujifilm Wako, 163-20145) for 45 min at 4°C. We removed the fixative, added 1 mL OWB, and stored samples at 4°C until staining. As an alternative, we assembled a D- or E-type chamber (CSCRIE) on a CREST glass slide (Matsunami Glass, SCRE-03) and directly loaded the organoid suspension into the chamber. In this format, we omitted centrifugation steps; all buffer exchanges were performed by manual pipetting, and the slide glass was kept overnight at 4°C in a humidified chamber.

Day 1: For the samples fixed in a tube, we spread organoids onto slide glass, maintained them in a humidified chamber at 4°C, and allowed them to adhere under their own weight for 1 h. We removed OWB and permeabilized organoids with 1x PBST at 4°C for 40 min. We removed PBST and incubated the organoids in OWB at 4°C for at least 15 min. We then removed OWB and incubated organoids with primary antibodies diluted 1:200 in OWB at 4°C overnight. The primary antibodies were rabbit anti-ZO-1 antibody (Proteintech, 21773-1-AP) and CD324 (E-cadherin) monoclonal antibody (DECMA-1, eBioscience, 14-3249-82).

Day 2: We washed organoids three times with OWB at room temperature, replacing the buffer every 2 h. We then incubated organoids with secondary antibodies (1:200) and 4′,6-diamidino-2-phenylindole (DAPI; final 0.5 µg/mL; Thermo Fisher, 2933025) in OWB at 4°C overnight. The secondary antibodies were goat anti-rabbit IgG (H+L) highly cross-adsorbed secondary antibody conjugated to Alexa Fluor Plus 647 (Thermo Fisher, A32733TR) and goat anti-rat IgG H&L secondary antibody conjugated to Alexa Fluor 488 (Abcam, ab150157).

Day 3: We washed organoids three times with OWB at room temperature, replacing the buffer every 2 h. We equilibrated ProLong Glass Antifade Mountant (Thermo Fisher, P36982) to room temperature for at least 1 h, replaced OWB with the mountant, and stored the samples at 4°C overnight.

Day 4: We returned the slides to room temperature and cured the mountant at 37°C for 1 h. We applied approximately 20 µL of glycerol (Fujifilm Wako, 072-00626), placed a coverslip on the sample, and sealed the edges with nail polish.

The images were recorded using a Stellaris 8 Laser Scanning Confocal Microscope (Leica Microsystems) with the following laser lines and detection windows: 405 nm (415 to 501 nm), 488 nm (503 to 654 nm), and 647 nm (657 to 831 nm).

### Image analysis

#### Overview and software environment

We performed all image analyses using custom Python scripts executed in Google Colab (Python 3.12.12). We implemented the analysis with standard Python libraries, including NumPy (ver. 2.0.2), pandas (ver. 2.2.2), SciPy (ver. 1.16.3), scikit-image (ver. 0.25.2), matplotlib (ver. 3.10.0), and seaborn (ver. 0.13.2). We processed microscopy images as TIFF files exported from an EVOS M7000 microscope and stored intermediate measurements as CSV or Excel files for downstream plotting and statistics.

### Common image-processing and segmentation workflow

Across analyses, we segmented target objects from fluorescence images using a consistent workflow. We first denoised the images using a median filter, followed by Gaussian smoothing (σ = 1 pixel). We then determined an intensity threshold to generate a binary mask (triangle or Otsu methods, as specified below). We refined binary masks by morphological operations (erosion and dilation using a 5 × 5 square structuring element) and hole filling. We labeled connected components and filtered objects based on predefined size and morphology criteria to exclude debris and artifacts. For area-to-length conversion, we converted pixel area to µm² using microscope calibration (µm per pixel). The equivalent circular diameter was calculated from the projected area of each segmented object by converting the measured area into the diameter of a circle with the same area. In this calculation, the radius corresponds to the square root of the projected area divided by pi, and the equivalent circular diameter is twice this radius.

### Quantification of Matrigel bead size

To quantify Matrigel bead size, we analyzed FITC fluorescence images of microbeads stained with FITC-lectin. We segmented microbeads using the common workflow described above, applying an Otsu threshold for binarization. Image fields containing obvious artifacts or segmentation failures were visually inspected and excluded from downstream analysis. We further excluded objects with diameters below 50 µm. Bead sizes were compared across conditions using box plots.

### Quantification of organoid size and number derived from organoid fragments

We quantified organoid size and number by segmenting AO-positive objects using the common workflow described above. For organoid size measurements, we analyzed non-tiled field images and excluded objects touching the image border. AO fluorescence images were binarized using an Otsu threshold, and objects with an equivalent circular diameter below 50 µm were excluded. We then inspected merged images of the segmentation labels overlaid on the corresponding phase contrast images and excluded obvious artifacts, partially segmented objects, single organoids split into multiple segments, and merged segments containing multiple organoids. This workflow was also used for the fragment size analysis in Figure 2. For downstream analyses, we defined organoids as AO-positive objects with an equivalent circular diameter of 100 µm or greater (∼7854 µm^2^).

For organoid counting, we analyzed stitched tiled images, counted organoids within a defined sample volume to obtain the suspension concentration, and then multiplied this value by the total suspension volume to estimate the total number of cultured organoids per sample. After automated image analysis, we removed obvious artifacts by reviewing segmentation overlays with the corresponding phase contrast images. Organoids were counted as objects exceeding the organoid definition threshold defined above.

### Quantification of organoid fragment size for passaging and post-passaging yield

For fragment quantification for passaging, AO images were segmented using triangle thresholding, and objects with a projected area of at least 5,000 µm² were counted.

We calculated the estimated total yield as follows: Estimated total yield = (Organoids from passaged fragments / Passaged fragments) × Total fragments collected.

### Quantification of crypt number and organoid formation rate

We quantified the number of crypts from tiled fluorescence images of PE-CD44 staining acquired across the entire imaging chamber. We segmented CD44-positive objects using triangle thresholding and excluded small objects with a projected-area cutoff of 2,500 µm².

After automated segmentation, we visually inspected overlays of the CD44 segmentation and corresponding phase contrast images to remove obvious artifacts. Crypt counts were obtained as the number of retained objects within the imaged chamber volume and converted to crypt concentration based on the known loaded volume.

In contrast to fragment-derived organoids, crypt-derived organoids are typically smaller on day 2. Accordingly, we defined crypt-derived organoids on day 2 as AO-positive objects with a projected area at least equivalent to a 50-µm-diameter circle (∼1,963.5 µm²). This cutoff was chosen based on phase contrast images overlaid with segmentation masks to exclude obvious debris and partial segments.

To quantify the organoid formation rate, we counted input crypts using CD44 image analysis as described above and quantified organoids on day 2 using AO staining and image analysis. We then calculated the formation rate as the number of day-2 organoids formed per input crypt.

### Statistics

Quantitative data are presented as mean ± SD unless otherwise stated. The number of independent experiments or analyzed objects is indicated in the corresponding figure legends. For object-level distributions, including Matrigel bead diameter and organoid projected area, each dot represents an individual segmented object, and these data were used to describe size distributions rather than to define independent biological replicates. For experiment-level comparisons, each dot represents one independent biological experiment.

Representative images are shown in Figures 2A, 2C, 2F, 3A, and 4A. Similar results were observed in at least three independent experiments unless otherwise indicated.

## Author contributions

Conceptualization, K.H. and H.K.; Methodology, H.K., K.H., M.M., and S.O.; Software, K.H.; Validation, H.K. and K.H.; Formal Analysis, K.H. and M.M.; Investigation, H.K. and K.H.; Resources, K.H.; Data Curation, K.H.; Writing-Original Draft, K.H.; Writing-Review & Editing, K.H., H.K., M.M., and S.O.; Visualization, K.H.; Supervision, K.H. and S.O.; Project Administration, K.H.; Funding Acquisition, K.H. and S.O.

## Data availability

Source data for all figures are available from the corresponding author upon reasonable request. The analysis code and representative raw images are available at https://github.com/solabtokyo-org/orb_2026.

## Acknowledgments

We thank all members of the Networked Biophotonics Group for their fruitful discussions. Part of Figure 1 was created with BioRender.com.

This work was supported by the Japan Agency for Medical Research and Development (AMED) under Grant Number JP22gm6710008 (to K.H.); the Japan Science and Technology Agency (JST) Fusion Oriented Research for disruptive Science and Technology (FOREST) Program (Grant Number JPMJFR240K to K.H.), Core Research for Evolutional Science and Technology (CREST) (Grant Numbers JPMJCR19H1 and JPMJCR23B6 to S.O.), Green Technologies of Excellence (GteX) Program (Grant Number JPMJGX23B1 to S.O.), and Adopting Sustainable Partnerships for Innovative Research Ecosystem (ASPIRE) program (Grant Numbers JPMJAP2416 and JPMJAP24B5 to S.O.); the Japan Society for the Promotion of Science (JSPS) Grants-in-Aid for Scientific Research (KAKENHI), including Grant-in-Aid for Challenging Research (Exploratory) (Grant Number JP25K22885 to K.H.) and Grant-in-Aid for Transformative Research Areas (A) (Grant Number JP25H01359 to S.O.); and the Nakatani Foundation (to S.O.), The Uehara Memorial Foundation (to K.H.), Takeda Science Foundation (to S.O. and K.H.), The Cell Science Research Foundation (to K.H.), Institute for Fermentation (to K.H.), Ono Medical Research Foundation (to K.H.), Nagase Science and Technology Foundation (to K.H.), and The Waksman Foundation of Japan (to K.H.).

